# Variation in Soil Properties under Long-Term Irrigated and Non-Irrigated Cropping and Other Land-Use Systems in Dura Catchment, Northern Ethiopia

**DOI:** 10.1101/755256

**Authors:** Gebreyesus Brhane Tesfahunegn, Teklebirhan Arefaine Gebru

## Abstract

There are limited reports about the impacts of long-term irrigated and non-irrigated cropping and land-use systems (CLUS) on soil properties and nutrient stocks under smallholder farmers’ conditions in developing countries. The objective of this research was to examine variation in soil properties and OC and TN stocks across the different CLUS in Dura sub-catchment, northern Ethiopia. Surveys and discussions on field history were used to identify nine CLUS, namely, tef *(Eragrostis tef* (Zucc) Trot)) mono-cropping (TM), maize *(Zea mays L*.) mono-cropping (MM), cauliflower (*Brassica oleracea var. botrytis*)-maize intercropping (IC1), red beet (*Beta Vulgaris*)-maize intercropping (IC2), cauliflower-tef-maize rotation (R1), onion (*Allium cepa* L.)-maize-onion rotation (R2), treated gully (TG), untreated gully (UTG), and natural forest system (NF). A total of 27 composite soil samples were collected randomly from the CLUS for laboratory analysis. Data were subjected to one-way analysis of variance and PCA. The lowest and highest bulk density was determined from NF (1.19 Mg m^-3^) and UTG (1.77 Mg m^-3^), respectively. Soil pH, EC and CEC varied significantly among the CLUS. The highest CEC (50.3 cmol_c_ kg^-1^) was under TG followed by NF. The highest soil OC stock (113.6 Mg C ha^-1^) and TN stock (12.2 Mg C ha^-1^) were found from NF. The PCA chosen soil properties explained 87% of the soil quality variability among the CLUS. Such soil properties and nutrient stocks variability among the CLUS suggested that introduction of suitable management practices are crucial for sustaining the soil system of the other CLUS.

## Introduction

Soil quality is becoming an important resource to raise crop productivity so that to meet the food required for the current and future population in developing countries as their economy mainly depends on agriculture [1-6]. Soil quality is defined as the capacity of the soil to give the intended functions for biomass and yield production [7-9]. In this study, the term soil quality is used synonymously with soil health. Recently, however, soil quality degradation caused by inappropriate cropping system, and land-use and soil management practices, has been reported among the top development challenges that demand urgent remedial actions. Several reports have shown that soil nutrient depletion and soil physical degradation are the dominant types of degradation associated with land and soil mis-management practices in the semi-arid areas [10-15].

Soil degradation poses serious development challenges in many developing countries including Ethiopia. Soil degradation induced by land and soil mis-management systems coupled with high dependency on erratic and unreliable rain-fed farming system has aggravated the problem of food insecurity in Ethiopia. Against such problem, irrigation agriculture has been suggested for implementing in the conditions of Ethiopia. For example, over 50 micro-dams were built-up since 1995, for being used mainly for irrigation purpose by smallholder farmers in the Tigray region, northern Ethiopia. Using the micro-dams in the region, so far a great deal of efforts has been attempted to achieve on sustainable economic, social and ecological developments. However, the sustainability of economic and ecological benefits from the micro-dams has been challenged by anthropogenic factors that increase sedimentation, soil and nutrients loadings and lowering of water use efficiency. Such factors have aggravated the rates of siltation of the micro-dams with less than 25% of their lifetime in Tigray region [16-19].

Even though there are problems of siltation and inefficient water use, irrigation agriculture from micro-dams as water source is becoming an essential government strategy for maximizing crop production per unit land area in Ethiopia conditions [20, 21]. Irrigation is designed to increase soil water availability and thereby enhances biomass production. The biomass is partly expected to be returned to the soil system to improve soil organic carbon (OC) and total nitrogen (TN) concentrations. Other practices such as reforestation of protected landscape, and agroforestry in agricultural lands have practiced to increase soil organic matter and soil nutrients for the past three decades in Ethiopia [9, 22-24]. Many researchers (e.g., Sharma et al. [4]; Lal [25]; Mandal et al. [26]; Yesilonis et al. [27]) have also reported that planting suitable crop types and cropping systems can play an important role in maintaining soil nutrients such as OC and TN stocks. However, increasing demand for short-term production encourages farmers to cultivate continuously (mono-cropping), overgrazed fields, or removed much of the above ground biomass through fuel collection, livestock feed and building materials. Eventually, such practices reduce soil nutrients and water holding capacity, increases erosion and thereby reduce agricultural productivity. For example, comparable higher OC, TN and other soil nutrients have reported under grassland as compared to cultivated land-use type [4, 14, 25, 28, 29]. However, there are limited quantitative evidences that evaluate soil properties variation under different cropping systems within the cultivated land-use type and compare with other land-use systems managed by smallholder farmers.

To maintain soil quality and reduce crop failures, intercropping which defined as the agricultural practice of cultivating two or more crops simultaneously in the same piece of land, has also reported by many researchers (e.g., Dallal [30]; Sharaiha et al. [31]; Nursima [32]). Intercropping is practiced commonly under irrigated agriculture and sometimes in rain-fed agriculture in northern Ethiopia even though their ecological benefits over the other cropping systems such as mono-cropping, rotation are not well documented (personal observation). The existing literatures have also shown that there is a need to understand impacts of continuous and other types of cropping systems on soil quality indicators in order to take appropriate measures that enhance sustainable crop production. The sustainability of soil for agricultural production can be viewed using soil properties as soil quality indicators (e.g., Arshad and Martin [2]; Trivedi et al. [14]; Iqbal et al. [28]; Andrews and Carroll [33]).

In developing countries such as Ethiopia, land has been utilized intensively for any purposes regardless of its suitability, which has resulted in severe soil quality degradation. Such degradation has explained by poor soil properties and low agricultural production [14, 34-38]. Practically, under the existing circumstances and economic conditions of farmers in developing countries such as Ethiopia there is a need to have inexpensive but environmentally sound integrated cropping system and land-use management approach to address soil quality related problems.

Increasing crop production in Ethiopia is likely to come from agricultural intensification and diversification through irrigation and other improved agronomic practices. Understanding impacts of different cropping and land-use practices on soil properties in general and soil OC and TN stock in particular is crucial for designing sustainable soil management practices. Scientific information on site-specific soil properties is a basic tool for proper soil management in order to provide sustainable soil functions at present and in the future [2, 4, 14, 25]. Site-specific data on soil properties could also support to deal with spatial variability of soil nutrients and physical indicators and their influencing factors. Such information is important to formulate appropriate sustainable cropping systems and land-use type strategies [6, 12, 14, 19].

The sustainability of crop and soil management practices to improve or maintain soil quality depends on understanding how soils respond to different site-specific cropping and land-use practices. Soil properties as indicators of soil functions and soil quality degradation status are suggested for understanding the sustainability of soil resources [2, 14, 39, 40]. There are many reports that generalized soil properties and soil nutrient stocks variability among different land-use types such as cultivated, grazing, grass and forest land (e.g., Trivedi et al. [14]; Chemeda et al. [15]; Wang et al. [19]; Fikadu et al. [24]; Lemenih and Hanna [41]; Batjes [42]; Yimer et al. [43]). However, there have been limited studies about the impacts of site-specific long-term irrigated and non-irrigated cropping and land-use systems on soil properties and soil nutrient stocks under smallholder farmers’ conditions in developing countries.

As most agricultural practices are site-specific the same management strategy cannot be said using the existing literature for the conditions of smallholder farmers in northern Ethiopia. Thus, it necessitates knowing the extent of soil quality degradation in terms of soil physical and chemical properties and nutrient stocks under different irrigation and rain-fed cropping and other land-use systems. There is also insufficient information about which soil properties (indicators) to be monitored over time with regard to the effects of cropping systems and other land-use practices in the study area conditions [5, 12, 19, 38]. This study was thus hypothesized that there is significant variability in soil properties and soil OC and TN stocks across the different long-term irrigated and non-irrigated adjacent cropping and land-use systems. The objectives of this research were to: (i) examine variation in soil properties under long-term irrigated and non-irrigated adjacent cropping and land-use systems (CLUS); (ii) evaluate soil organic carbon and total nitrogen stocks across the different CLUS; and (iii) examine soil properties that explain better for soil quality variability across the different CLUS in the conditions of Dura sub-catchment, northern Ethiopia.

## Materials and Methods

### 2.1 Study area

This research was carried-out from February 2015 to June 2015 in Dura sub-catchment of Tigray region, northern Ethiopia (Fig 1). The study catchment area covers about 5000 ha and area of its sub-catchment is 1240 ha. Altitude of the sub-catchment ranges between 2050 and 2650 m above sea level [44]. In the study sub-catchment, mean annual temperature of 22°C and rainfall of 700 mm were reported using 35 years of meteorological data. The study sub-catchment receives normal rainfall during June to early September which is unimodal (Meteorology Agency-Mekelle branch). Crop and livestock mixed-farming is commonly practiced. However, crop production is the dominant farming system for farmers’ livelihood. Arable land is dominated over the other land-use types in the study catchment.

**Figure 1.**
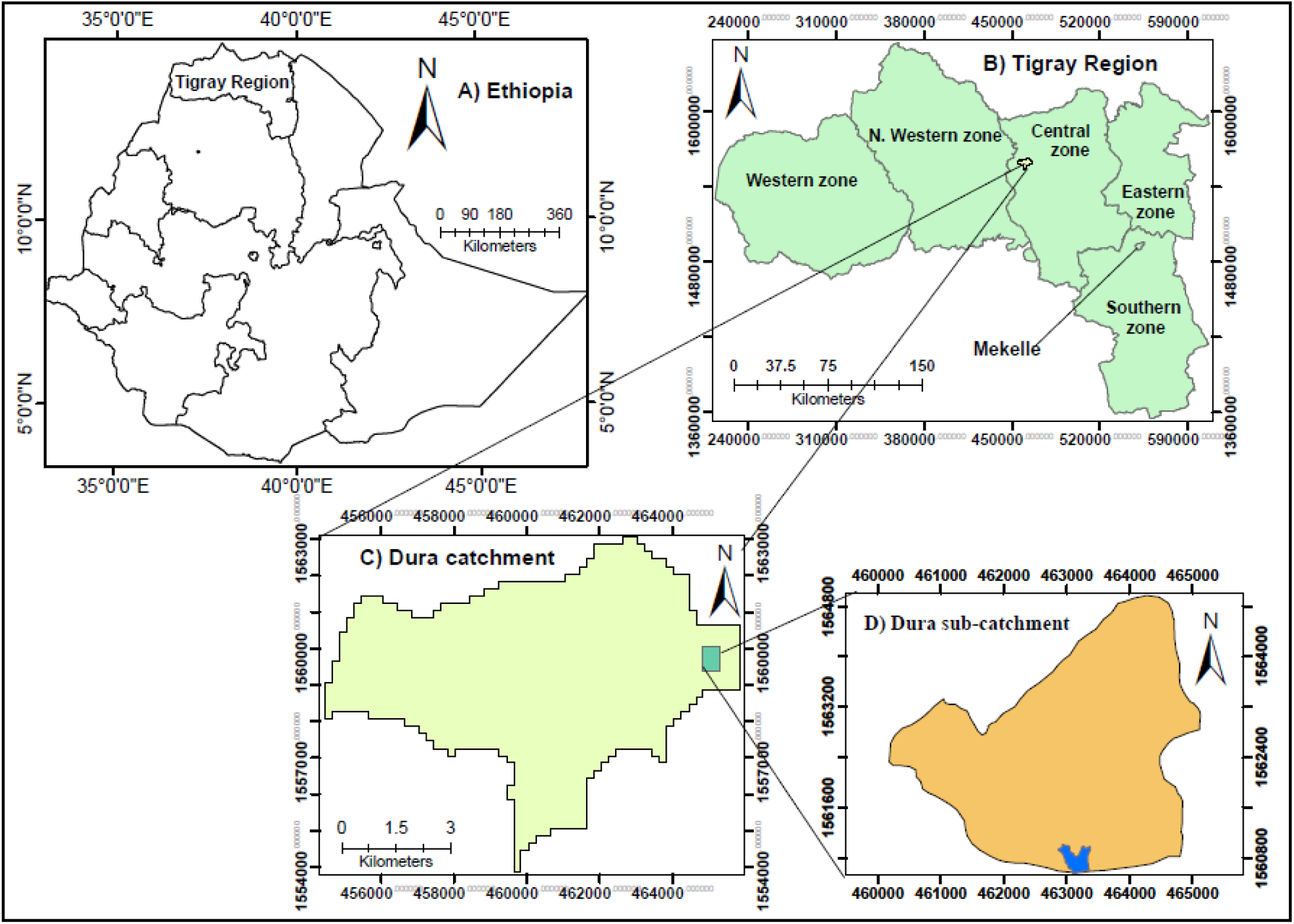
Location of the study area: Ethiopia (A), Tigray Region (B), Dura catchment (C) and Dura sub-catchment (D). Blue color in the sub-catchment is reservoir.

In Dura sub-catchment, both rain-fed and irrigation agriculture have been practiced for more than 2 decades. But rain-fed agriculture which is the oldest practice dominated in area coverage. About 100 ha farmland has been irrigated since 1996 in Dura sub-catchment. Afforested area, pasture, scattered woody trees, bushes and shrub lands were also found in the study sub-catchment. The dominant soils in the study sub-catchment includes: Eutric Cambisols on the steep slope, Chromic Cambisols on the middle to steep slopes and Chromic Vertisols on the flat areas [45]. This study sub-catchment was selected as it represents the mid-highland agro-ecology conditions with practices of long-term irrigated and non-irrigated adjacent fields under smallholder farmers in northern Ethiopia.

### 2.2 Identification of long-term irrigated and non-irrigated cropping and land-use systems

Reconnaissance surveys coupled with formal and informal group discussions with farmers and development agents (DAs) were used to identify the different irrigated and non-irrigated cropping and land-use systems in Dura sub-catchment, northern Ethiopia. During the field surveys in February and March 2015 the researcher together with the DAs visited the study catchment to get an overall impression about the irrigation command area, adjacent rain-fed cropping and land-use systems. The purpose of the reconnaissance survey was to characterize the fields’ historical cropping system, soil management, agronomic practices and field features. During the survey, participatory tools such as field observation, transect-walks and group discussion were employed. The transect-walks were done twice, that is, from the east to the west and also from the north to south direction of the study sub-catchment in order to observe different cropping and land-use systems. This was done by the team composed of the researcher, three (3) DAs and randomly selected 10 farmers from the study sub-catchment.

Three group discussions sessions were held in order to reach consensus among the participants about the descriptions of the irrigated and non-irrigated fields that were selected. On the basis of the farmers’ final consensus nine (9) dominant cropping and land-use systems (fields) were identified (Table 1), and geo-referenced and described their topographic features (Table 2). Such fields were selected because the site and crop specific management practices perhaps affect the sustainability of natural resources, crop productivity and soil fertility utilization in the catchment. The selected fields were located on Chromic Vertisols adjacent to each other at a distance that ranges between 50 and 150 m.

**Table 1.**
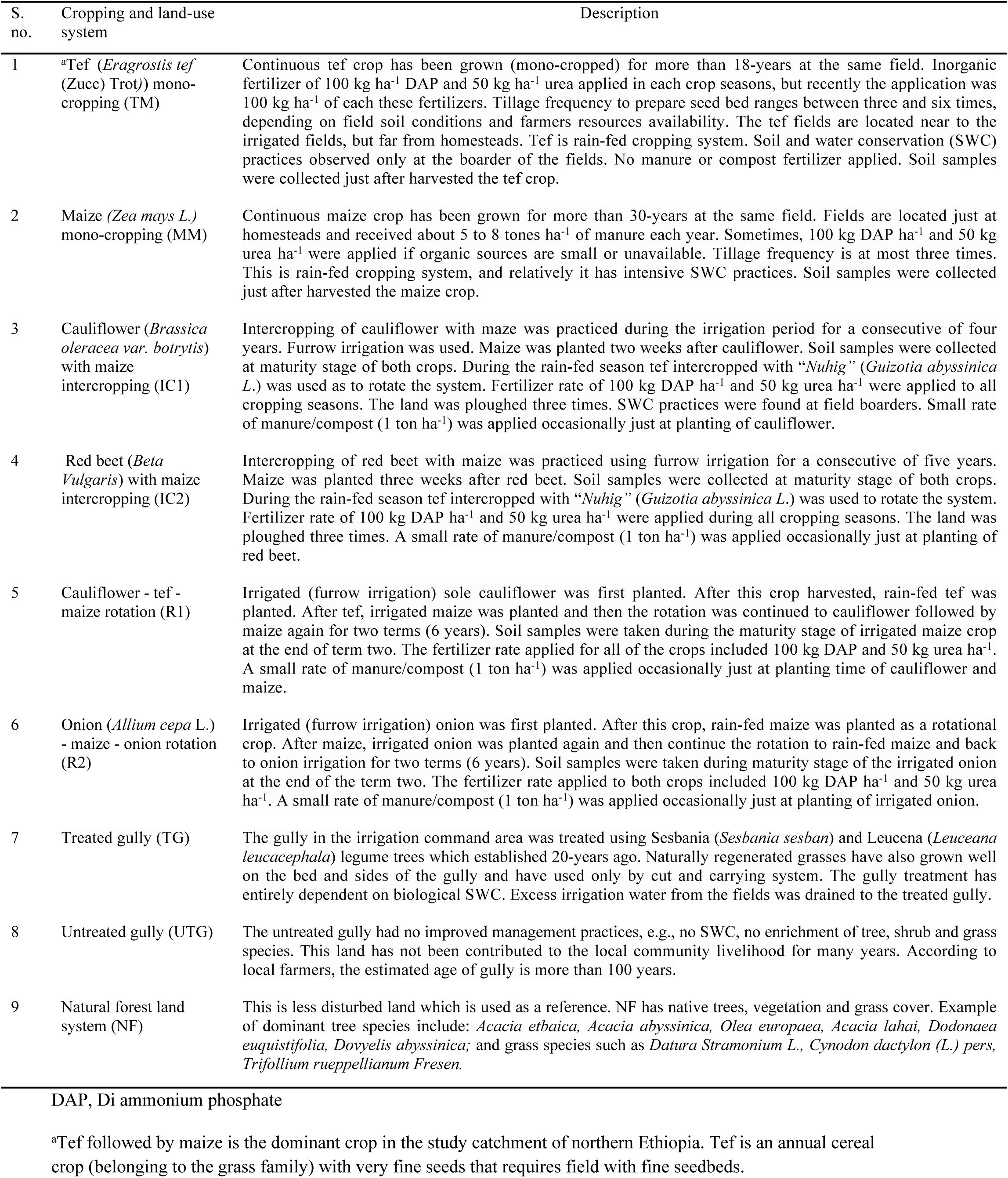
Cropping and land-use systems identified in the Dura sub-catchment, northern Ethiopia.

| S. no. | Cropping and land-use system | Description |
| --- | --- | --- |
| 1 | <sup>a</sup>Tef (*Eragrostis tef* (Zucc) Trot)) mono-cropping (TM) | Continuous tef crop has been grown (mono-cropped) for more than 18-years at the same field. Inorganic fertilizer of 100 kg ha<sup>-1</sup> DAP and 50 kg ha<sup>-1</sup> urea applied in each crop seasons, but recently the application was 100 kg ha<sup>-1</sup> of each these fertilizers. Tillage frequency to prepare seed bed ranges between three and six times, depending on field soil conditions and farmers resources availability. The tef fields are located near to the irrigated fields, but far from homesteads. Tef is rain-fed cropping system. Soil and water conservation (SWC) practices observed only at the boarder of the fields. No manure or compost fertilizer applied. Soil samples were collected just after harvested the tef crop. |
| 2 | Maize (*Zea mays* L.) mono-cropping (MM) | Continuous maize crop has been grown for more than 30-years at the same field. Fields are located just at homesteads and received about 5 to 8 tones ha<sup>-1</sup> of manure each year. Sometimes, 100 kg DAP ha<sup>-1</sup> and 50 kg urea ha<sup>-1</sup> were applied if organic sources are small or unavailable. Tillage frequency is at most three times. This is rain-fed cropping system, and relatively it has intensive SWC practices. Soil samples were collected just after harvested the maize crop. |
| 3 | Cauliflower (*Brassica oleracea* var. *botrytis*) with maize intercropping (IC1) | Intercropping of cauliflower with maze was practiced during the irrigation period for a consecutive of four years. Furrow irrigation was used. Maize was planted two weeks after cauliflower. Soil samples were collected at maturity stage of both crops. During the rain-fed season tef intercropped with “*Nuhig*” (*Guizotia abyssinica* L.) was used as to rotate the system. Fertilizer rate of 100 kg DAP ha<sup>-1</sup> and 50 kg urea ha<sup>-1</sup> were applied to all cropping seasons. The land was ploughed three times. SWC practices were found at field boarders. Small rate of manure/compost (1 ton ha<sup>-1</sup>) was applied occasionally just at planting of cauliflower. |
| 4 | Red beet (*Beta Vulgaris*) with maize intercropping (IC2) | Intercropping of red beet with maize was practiced using furrow irrigation for a consecutive of five years. Maize was planted three weeks after red beet. Soil samples were collected at maturity stage of both crops. During the rain-fed season tef intercropped with “*Nuhig*” (*Guizotia abyssinica* L.) was used to rotate the system. Fertilizer rate of 100 kg DAP ha<sup>-1</sup> and 50 kg urea ha<sup>-1</sup> were applied during all cropping seasons. The land was ploughed three times. A small rate of manure/compost (1 ton ha<sup>-1</sup>) was applied occasionally just at planting of red beet. |
| 5 | Cauliflower - tef - maize rotation (R1) | Irrigated (furrow irrigation) sole cauliflower was first planted. After this crop harvested, rain-fed tef was planted. After tef, irrigated maize was planted and then the rotation was continued to cauliflower followed by maize again for two terms (6 years). Soil samples were taken during the maturity stage of irrigated maize crop at the end of term two. The fertilizer rate applied for all of the crops included 100 kg DAP and 50 kg urea ha<sup>-1</sup>. A small rate of manure/compost (1 ton ha<sup>-1</sup>) was applied occasionally just at planting time of cauliflower and maize. |
| 6 | Onion (*Allium cepa* L.) - maize - onion rotation (R2) | Irrigated (furrow irrigation) onion was first planted. After this crop, rain-fed maize was planted as a rotational crop. After maize, irrigated onion was planted again and then continue the rotation to rain-fed maize and back to onion irrigation for two terms (6 years). Soil samples were taken during maturity stage of the irrigated onion at the end of the term two. The fertilizer rate applied to both crops included 100 kg DAP ha<sup>-1</sup> and 50 kg urea ha<sup>-1</sup>. A small rate of manure/compost (1 ton ha<sup>-1</sup>) was applied occasionally just at planting of irrigated onion. |
| 7 | Treated gully (TG) | The gully in the irrigation command area was treated using Sesbania (*Sesbania sesban*) and Leucena (*Leuceana leucacephala*) legume trees which established 20-years ago. Naturally regenerated grasses have also grown well on the bed and sides of the gully and have used only by cut and carrying system. The gully treatment has entirely dependent on biological SWC. Excess irrigation water from the fields was drained to the treated gully. |
| 8 | Untreated gully (UTG) | The untreated gully had no improved management practices, e.g., no SWC, no enrichment of tree, shrub and grass species. This land has not been contributed to the local community livelihood for many years. According to local farmers, the estimated age of gully is more than 100 years. |
| 9 | Natural forest land system (NF) | This is less disturbed land which is used as a reference. NF has native trees, vegetation and grass cover. Example of dominant tree species include: *Acacia etbaica*, *Acacia abyssinica*, *Olea europaea*, *Acacia lahai*, *Dodonaea euquistifolia*, *Dovyelis abyssinica*; and grass species such as *Datura Stramonium* L., *Cynodon dactylon* (L.) pers, *Trifolium rueppellianum* Fresen. |

DAP, Di ammonium phosphate
<sup>a</sup>Tef followed by maize is the dominant crop in the study catchment of northern Ethiopia. Tef is an annual cereal crop (belonging to the grass family) with very fine seeds that requires field with fine seedbeds.

**Table 2.**
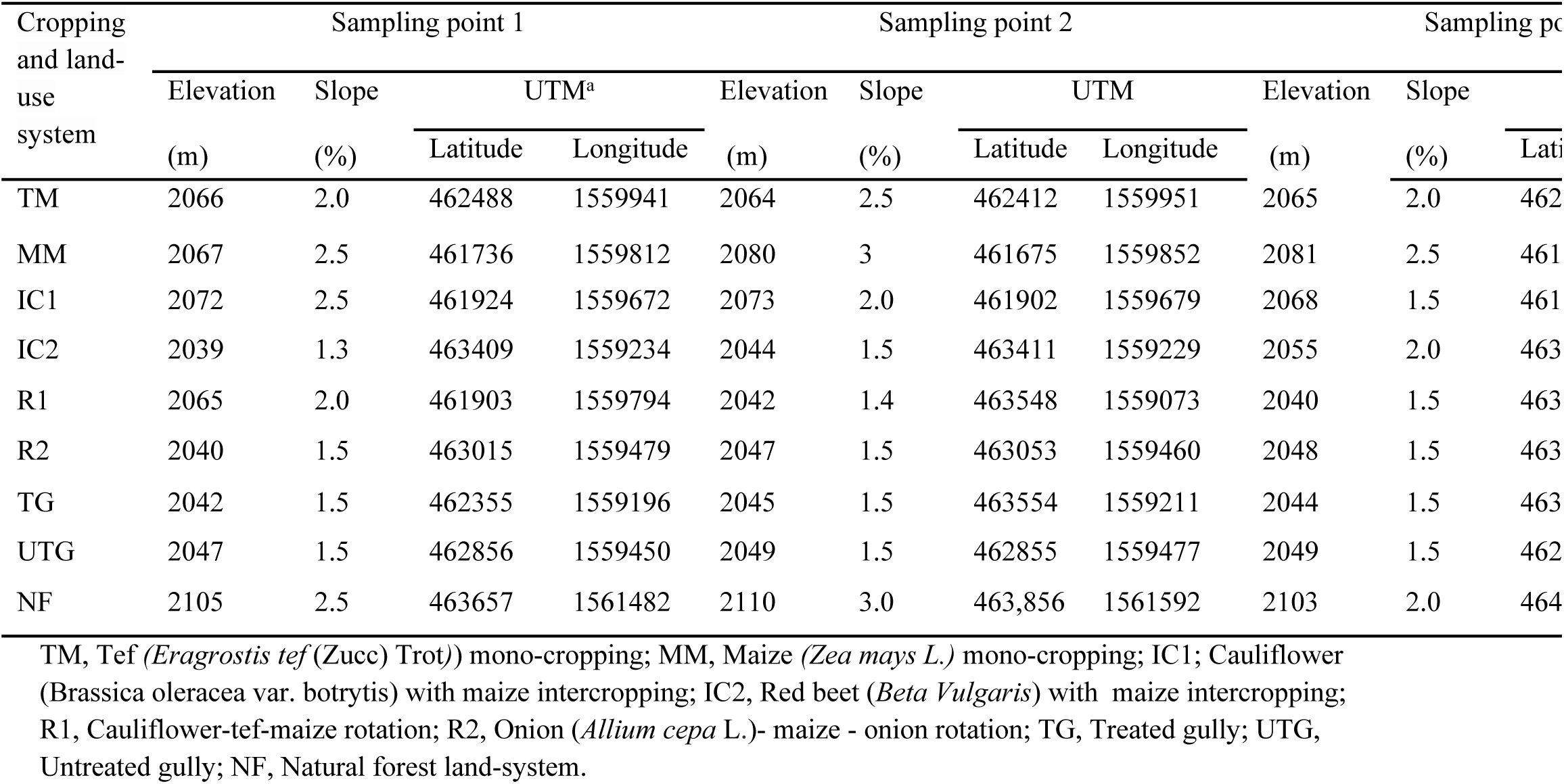
Topographic features of each soil sampling unit selected from the long-term irrigated and non-irrigated cropping and land-use systems in the Dura sub-catchment, northern Ethiopia.

From the total nine (9) fields identified, four (4) were from irrigated fields, two (2) from rain-fed cropping system, and three (3) from other land-use systems. The cropping and land-use systems (fields) selected were: (i) Tef *(Eragrostic tef* (Zucc) Trot)) mono-cropping (TM), (ii) Maize *(Zea mays L*.) mono-cropping (MM), (iii) Cauliflower (*Brassica oleracea* var. botrytis) with maize intercropping (IC1), (iv) Red beet (*Beta Vulgaris*) with maize intercropping (IC2), (v) Cauliflower - tef - maize rotation (R1), (vi) Onion (*Allium cepa* L.) - maize - onion rotation (R2), (vii) Treated gully (TG), (viii) Untreated gully (UTG), and (ix) Natural forest land-use system (NF) (Table 1). The first two (TM and MM) were selected from the rain-fed fields adjacent to the irrigation command area, whereas IC1, IC2, R1 and R2 were selected from the irrigated crop fields. TG and UTG were also found within the irrigation command area. An adjacent natural forest land-use system (NF) was used as a reference while compared with the impact of irrigation and non-irrigation cropping and land-use systems on soil properties, and carbon and nitrogen stocks.

### 2.3 Sample designing, soil sampling and analysis

The targeted fields (population) were all the cropping and land-use systems (CLUS) practiced in Dura sub-catchment. Farmers were selected nine CLUS that dominantly available in the sub-catchment. The soil samples were taken from the nine irrigated and non-irrigated CLUS which were selected from the sub-catchment. Composite soil samples were collected from the selected sampling points in each CLUS using judgmental sampling on the basis of reliable historical and physical knowledge of experts and local farmers (Table 1). Soil samples were collected in May 2015. Three sampling units replicated in the nine CLUS were identified by experts’ judgment based on field homogeneity. Identification of the sampling units using expert knowledge is very efficient as it is quick and easy to select the sampling units.

Considering the costs of soil analysis and its statistical representativeness a total of 27 composite soil samples from three sampling units (9 CLUS × 3 sampling units) were collected. The soil sampling unit plot area was 48 m^2^ (6 m × 8 m). The soil sampling plots land features are described in Table 2. In each sampling unit plot 10 pairs of randomly selected coordinate points were identified. From the 10 geo-referenced points in each sampling plot, three sampling points were selected using simple random sampling technique whereby the composite soil sample from each plot was collected.

The soil samples were taken from each sampling point at 0-20 cm soil depth. This sampling depth is where most soil changes are occurred due to long-term cropping systems, land-use types, and soil and water management practices including irrigation agriculture. Three soil samples were collected from each sampling unit in a plot and pooled (composited) into a bucket and mixed thoroughly to form a single homogenized sample. A sub-sample of 500 g soil that represented the pooled sample in the bucket was taken from each sampling unit plot, and air dried and sieved through 2 mm mesh sieves. In addition, three undisturbed soil samples were collected from each irrigated and non-irrigated field sampling unit plots at 0-20 cm soil depth using 5·0 cm long by 5·0 cm diameter cylindrical metal core sampler to determine soil dry bulk density.

The analysis of the soil samples was carried-out following the standard laboratory procedures in JIJE Analytical Testing Service Laboratory in Addis Ababa, Ethiopia. Soil texture was determined using the Bouyoucos hydrometer method [46] and soil dry bulk density (DBD) by the core method [47]. Total porosity was calculated from the DBD and assumed average particle density (PD) of 2.65 Mg m^-3^ as (1-DBD/PD) × 100 [48]. The depth of A-horizon was directly measured as an average of the three pits opened (0·60 m depth) in each sampling unit plot.

Soil pH was determined in 1:2·5 soil to water ratio using pH meter combined glass electrode [49], electrical conductivity (EC) in 1:2·5 soil to water ratio using conductivity meter [50], soil organic carbon (OC) by the Walkley-Black method [51], available phosphorus (Pav) by Olsen method [52], and total nitrogen (TN) by Kjeldhal digestion method followed by distillation and titration [53]. Cation exchange capacity (CEC) was determined by ammonium acetate extraction buffered at pH 7 using flame photometer [54].

Exchangeable bases (Ca^2+^, Mg^2+^, K^+^ and Na^+^) were analyzed after extracted in a 1:10 soil/solution ratio using 1M ammonium acetate at pH 7.0. Readings for Ca^2+^ and Mg^2+^ in the extracts were analyzed using atomic absorption spectrophotometer whereas Na^+^ and K^+^ were determined by flame photometry [55].

### 2.4 Derived other soil parameters

Soil structural stability index is estimated [56, 57] as:

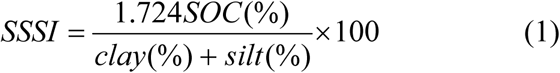

Where, SSSI (%) is soil structural stability index, SOC is soil organic carbon, and clay + silt is combined clay and silt content. SSSI < 5% indicates structurally degraded soil; 5% < SSSI < 7% indicates high risk of soil structural degradation; 7% < SSSI < 9 % indicates low risk of soil structural degradation; and SSSI > 9% indicates sufficient SOC to maintain the structural stability. A higher the SSI value, a better would be in maintaining soil structural degradation.

Base saturation percentage was calculated by divided the sum of base forming cations (Ca^2+^, Mg^2+^, K^+^ and Na^+^) by CEC and then multiplied by 100%. Exchangeable sodium percentage (ESP) was calculated by divided exchangeable Na^+^ by CEC. The ESP threshold of 15% was used to classify sodium hazard, that is, sodic soils are those with ESP of more than 15%. Sodium adsorption ratio (SAR) was calculated [58-60] as:

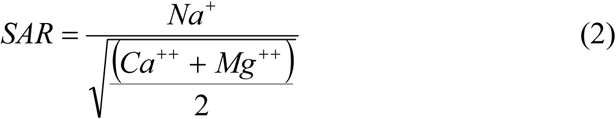

Where, SAR is sodium adsorption ratio in (cmol kg^-1^)^0.5^; and Na^+^, Mg^2+^ and Ca^2+^ are exchangeable sodium, magnesium and calcium, respectively, in cmol_c_ kg^-1^. SAR < 12 indicate non sodicity and values >12 indicate sodic soils [58].

The relationship between soil OC and TN as represented by the ratio of OC to TN was derived as an indicator of soil quality status. The OC: TN is a sensitive indicator of soil quality when assesses soil carbon and nitrogen nutrient balance. It is used as a sign of soil nitrogen mineralization capacity [61, 62]. A high OC: TN indicates the slowdown decomposition rate of organic matter by limiting soil microbial activity. On the other hand, low ratio of OC: TN could show the accelerated process of microbial decomposition on organic matter and nitrogen, in which this is not conducive for carbon sequestration [61-63].

Soil OC and TN stocks (Mg ha^-1^) were calculated using the model developed by Ellert and Bettany [64] as:

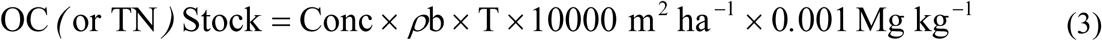

Where: *OC (or TN) Stock* is soil organic carbon or total nitrogen stock (Mg ha^-1^), *Conc* is soil organic carbon or total nitrogen concentration (kg Mg^-1^), *ρb* is dry bulk density (Mg m^-3^) and *T* is thickness of soil layer (m). Natural forest (less disturbed system) was used as a reference while assessed the amount of soil OC and TN stock reduction due to the effects of each CLUS. Thus, the difference between NF and any CLUS divided by NF and multiplied by 100% was used to show the reduction of OC and TN from the soil stocks in the different CLUS. Reduction in soil OC and TN implies there is contribution towards increasing the global green house gases that emitted to the atmosphere.

### 2.5 Data analysis

Data were analyzed using statistical software package of SPSS 20·0, SPSS Inc. International Business Machines Company, Chicago, USA. One-way analysis of variance was carried out to test the mean differences of the soil properties among the nine cropping and land-use systems. Data were tested for the assumption of normal distribution. Means were separated using Least Significant Differences at probability level (P) ≤ 0·05. Data were also subjected to descriptive, correlation (r) and factor analysis.

Correlations among the soil properties were checked by Pearson product moment correlation test (two-tailed) at P ≤ 0·05 in order to assess the effect of multi-collinearity. The principal component analysis (PCA) was also used to extract factor components and reduce variable redundancy. The PCA was thus used to examine the relationship among the 22 soil properties by statistically grouped into five principal components (PCs) using the Varimax rotation procedure. The five PCs with eigenvalues > 1 that explained at least 5% of the variation of the soil properties response to the cropping and land-use systems were considered. Varimax rotation with Kaiser Normalization resulted in a factor pattern that highly loads into one factor [65]. If the highly weighted variables within PC correlated at the correlation (r) value < 0.60, all variables were retained in the PC. Among the well-correlated variables (r ≥ 0.60) within PC, a variable with the highest partial correlation coefficient and factor loading was retained in the component factor. Note that only variables with factor loadings > 0.7 were retained in the PC. If the loading coefficient of a variable was > 0.7 in more than one component, it was suggested to select from the component holding with the highest coefficient for that variable [66]. Communalities that estimate the portion of variance of each soil parameter in the component factor was also considered while selected a variable to be retained in the PC. A higher communality for a soil parameter indicates a higher proportion of the variance is explained the component factor by the variable. Less importance should be ascribed to soil parameters with low communalities when interpreting the PC factors [65].

### 2.6 Ethics approval

Ethical approval was obtained from the Research and Community Services Directorate Director Research Ethics Review Committee of Aksum University, Ethiopia to conduct this study. Full right was given to the study participants to refuse and withdraw from participation at any time. Confidentiality of respondents was preserved by the researchers during data collection of soil samples and soil and crop management history. It was also noted that the research has no any activities that directly related to human being as it is directly related to the physical environment.

## 3. Results ad Discussion

### 3.1 Effect on soil physical properties

The soil physical properties significantly varied among most cropping and land-use systems (CLUS) in Dura sub-catchment, northern Ethiopia. There were significant differences in clay, silt, sand, bulk density, porosity, soil structural stability index, and A-horizon among most of the CLUS (Table 3). The soil clay contents of the CLUS varied significantly between 26 to 74%, with the lowest and the highest values observed from TM and R2, respectively. This could influence the other textural classes and physical and chemical soil properties.

**Table 3.** Mean soil physical properties variability among the cropping and land-use systems at 0-20 cm depth in the Dura sub-catchment, northern Ethiopia.

| Physical soil property | Cropping and land-use system (CLUS) |  |  |  |  |  |  |  |  | Mean |
| --- | --- | --- | --- | --- | --- | --- | --- | --- | --- | --- |
|  | TM | MM | IC1 | IC2 | R1 | R2 | TG | UTG | NF |  |
| Clay (%) | 26.3f | 32.0e | 43.7d | 48.7c | 54.0b | 73.7a | 49.7c | 36.7e | 33.7e | 44.2 |
| Silt (%) | 33.4d | 39.0b | 36.7c | 30.0e | 33.7d | 22.7f | 34.7cd | 25.3f | 43.3a | 33.3 |
| Sand (%) | 40.3a | 29.0b | 19.7d | 21.3d | 12.3e | 3.7f | 15.7e | 38.0a | 23.0c | 22.6 |
| DBD (Mg m<sup>-3</sup>) | 1.59b | 1.39f | 1.47cd | 1.50c | 1.43e | 1.45de | 1.32g | 1.77a | 1.19h | 1.46 |
| TP (%) | 40.1g | 47.5c | 44.5ef | 43.4f | 46.2d | 45.3de | 50.3b | 33.2h | 55.1a | 45.1 |
| SSSI (%) | 1.85f | 3.93c | 2.68d | 2.67d | 2.218e | 1.99e | 7.17b | 1.15g | 12.1a | 3.89 |
| A-h (m) | 0.104d | 0.138c | 0.110d | 0.102d | 0.105d | 0.106d | 0.187b | 0.017e | 0.195a | 0.120 |

Means followed by different letters in the same row are significantly different at probability level (P) = 0.05.
DBD, dry bulk density; TP, total porosity; SSSI, soil structural stability index; A-h, A-horizon depth; TM, Tef(*Eragrostis tef* (Zucc) Trot)) mono-cropping system; MM, Maize (*Zea mays L.*) mono-cropping system; IC1; Cauliflower (*Brassica oleracea* var. botrytis) with maize intercropping; IC2, Red beet (*Beta Vulgaris*) with maize intercropping; R1, Cauliflower - tef - maize rotation; R2, Onion (*Allium cepa L.*) - maize - onion rotation; TG, Treated gully; UTG, Untreated gully; NF, Natural forest.

The lowest silt (22.7%) and sand (3.7%) contents were observed from R2 whereas the highest silt (43%) from NF and sand (39.0%) from TM were observed. The highest sand content in the TM may be associated with repeated cultivation using inorganic fertilizer for long-time in which such practices aggravate erosion that erode fine soil particles and leaves coarser particles (Brady and Weil, 2002). The mean clay (44.2%), silt (33.2%) and sand (22.6%) contents of all the CLUS indicated that the sub-catchment soil has dominated by clay followed by silt texture soil. Fields with higher clay content such as R2 are considered by local farmers as difficult for workability and so susceptible to the problem of water logging in which this is in agreement with the reports reported by Barrios and Trejo [67]; Mairura et al. [68] and Tesfahunegn et al. [69].

On the other hand, there were non-significant differences in soil sand contents among some of the CLUS, e.g., between TM and UTG, IC1 and IC2, R1 and TG (Table 3). This could be attributed to the fact that sand texture is soil property that does influence little by some of the CLUS and their activities and so by erosion-deposition processes. The present finding on soil sand content is consistent with Shepherd et al. [70] who reported that there is no significant effect of land-use systems on soil particle size distribution. However, such reports contradicted with that of Kauffmann et al. [71]; Voundi Nkana and Tonye [72] and Agoumé and Birang [73] who reported that continuous cropping and intensive land-use systems have significantly affected soil particle size distribution. Such discrepancy of results on soil particles could be attributed to the duration of the cropping system, variability of management practices and weather conditions, and effects of variation in topography.

The lowest dry bulk density (DBD) was recorded from NF (1.19 Mg m^-3^) followed by TG (1.32 Mg m^-3^) and MM (1.39 Mg m^-3^). Conversely, the highest bulk density was found from UTG (1.77 Mg m^-3^) followed by TM (1.59 Mg m^-3^) and IC2 (1.50 Mg m^-3^). However, there were non-significant differences in DBD between IC1 and IC2, and R1 and R2. Generally, DBD was found to be higher in mono-cropping than intercropping and rotation cropping systems; and in intercropping than crop rotation fields (Table 3). The exceptional lower DBD from MM could be associated with the effect of manure and compost whereby farmers regularly applied to maize fields in each cropping season. The DBD of NF, TG and MM were found within the ideal critical levels (1·00-1·40 Mg m^-3^) as described as an ideal soil condition for plant root growth and water holding capacity by Arshad et al.[74]. However, the other cropping and land-use systems considered in this study showed DBD values higher than the critical level in which this implies the need for adopting appropriate practices that improve soil bulk density.

Total porosity, SSSI and A-horizon values were significantly varied among the CLUS, with the highest of these parameters recorded from NF and the lowest from UTG (Table 3). The trend of these parameters is similar to that of DBD but in the opposite direction. The variation in total porosity, SSSI and A-horizon among the different CLUS could be attributed to the differences in soil organic matter (SOM) contents. In fact, land-use systems such as NF, TG and MM which have received higher OM sources can improve the quality of soil properties. Soils with a good physical quality have a stable structure which resist for the effects of erosion [73, 74]. The risk of soil structural degradation associated with SOC depletion was found to be higher in UTG followed by TM even though R1, R2, IC1, IC2 and MM are also showed structurally degraded soil with SSSI < 5%.

Consistent with the present finding, substantial reports have shown that degraded soil that receives higher SOM can improve soil porosity, soil structure and depth of A-horizon. Improving such soil attributes would enhance soil water-holding capacity, decreases runoff and soil losses and eventually increases agricultural production (e.g., Agoumé and Birang [73]; Arshad et al. [74]; Sojka and Upchurch [75]; Evanylo and McGuinn [76]; Sally and Karle [77]; Moghadam et al. [78]). Conversely, cultivated fields treated with mineral fertilizer consecutively for many years such as TM and irrigation fields (e.g., IC1, IC2) showed poor soil physical properties. The implication of this study result is that cropping systems treated using only mineral fertilizer for a long-term deteriorates soil physical properties. However, the trends of rates of mineral fertilizer have been increased from time to time in Ethiopia. The present result of structural physical degraded soils could be associated with the continuous application of chemical fertilizer for more than 2 decades which in line with the reports from Moghadam et al. [78] and Ayoola [79] who reported a negative effect of continuous usage of mineral fertilizer on the soil system.

### 3.2 Effect on soil chemical properties

#### 3.2.1 Effects on soil pH and electrical conductivity

There were statistically significant differences in soil chemical properties among most cropping and land-use systems (CLUS) in the study sub-catchment (Table 4). The soil pH varied significantly from 6.94 in TM to 8.50 in R1. The higher pH in R1 could be associated with the effects of long-term irrigation and soil management practices. There was also non-significant differences in soil pH among some of the CLUS (e.g., between MM and NF; and among IC1, IC2, and TG). The mean soil pH (7.68) of all the CLUS indicates that the study catchment soil is categorized as moderately alkaline in reference to the classification for African soils reported by Landon [80]. Generally, the CLUS in the catchment showed soil pH values within the critical levels (6.5-8.5) reported in literature (e.g., Sanchez et al. [81]; Tesfahunegn et al. [82], which indicates that soil pH is not a key problem to monitor effects of the different cropping and land-use systems on soil quality indicators.

**Table 4.** Mean soil chemical properties variability among the cropping and land-use systems at 0-20 cm depth in the Dura sub-catchment, northern Ethiopia.

| Chemical soil property | Cropping and land-use system (CLUS) |  |  |  |  |  |  |  |  |  |
| --- | --- | --- | --- | --- | --- | --- | --- | --- | --- | --- |
|  | TM | MM | IC1 | IC2 | R1 | R2 | TG | UTG | NF | Mean |
| Soil pH | 6.94f | 7.31de | 7.98bc | 8.35ab | 8.50a | 7.79c | 7.91bc | 7.09f | 7.28e | 7.68 |
| EC (ds m<sup>-1</sup>) | 0.057c | 0.333ab | 0.327ab | 0.510a | 0.273b | 0.390ab | 0.267b | 0.210bc | 0.230bc | 0.289 |
| CEC (cmol<sub>c</sub> kg<sup>-1</sup>) | 18.8f | 37.8b | 28.9e | 30.4de | 32.2cd | 34.0c | 50.3a | 11.4g | 49.8a | 32.6 |
| Ex Ca (cmol<sub>c</sub> kg<sup>-1</sup>) | 8.5f | 18.3b | 12.1e | 13.3d | 14.0cd | 14.8c | 24.8a | 4.5g | 25.5a | 15.1 |
| Ex Mg (cmol<sub>c</sub> kg<sup>-1</sup>) | 2.7f | 8.8c | 6.3e | 6.5de | 7.1d | 6.8d | 14.7b | 1.3g | 15.1a | 7.6 |
| Ex Na (cmol<sub>c</sub> kg<sup>-1</sup>) | 0.040f | 0.213d | 0.667a | 0.682a | 0.627b | 0.613b | 0.450c | 0.153e | 0.030f | 0.386 |
| Ex K (cmol<sub>c</sub> kg<sup>-1</sup>) | 0.503d | 0.797b | 0.605c | 0.597c | 0.623c | 0.627c | 1.43a | 0.204e | 1.39a | 0.753 |
| SBFC (cmol<sub>c</sub> kg<sup>-1</sup>) | 11.7e | 28.1b | 19.7d | 21.1cd | 22.4c | 22.9c | 41.4a | 6.2f | 42.0a | 23.9 |
| ESP | 0.12f | 0.45 | 1.40a | 1.26b | 1.21b | 0.97c | 0.84d | 0.28e | 0.06f | 0.73 |
| BSP | 62.5d | 74.3b | 68.2cd | 69.4b | 69.6b | 67.4d | 82.3a | 54.4e | 84.3a | 73.3 |
| SAR | 0.011f | 0.049 | 0.153a | 0.140ab | 0.139b | 0.117c | 0.098d | 0.034e | 0.007f | 0.083 |
| Pav (mg kg<sup>-1</sup>) | 2.0gh | 13.8c | 2.8e | 3.1de | 3.5d | 4.0d | 19.4b | 1.4h | 23.9a | 8.2 |
| OC (%) | 0.643f | 1.540c | 1.250d | 1.220d | 1.110e | 1.113e | 3.520b | 0.413g | 4.980a | 1.754 |
| TN (%) | 0.067f | 0.135c | 0.119d | 0.116d | 0.108e | 0.109e | 0.257b | 0.034g | 0.385a | 0.148 |
| OC:TN | 9.597g | 11.407d | 10.50e | 10.51e | 10.28f | 10.21f | 13.696a | 12.147c | 12.935b | 11.87 |

Means followed by different letters in the same rows are significantly different at probability level (P) = 0.05.
pH, hydrogen ion concentration; EC, Electrical conductivity; CEC, cation exchange capacity; Ex Ca, Ex Mg, Ex Na, Ex K, exchangeable calcium, magnesium, sodium, potassium, respectively; SBFC sum of base forming cations; ESP, Exchangeable Na percentage; BSP, Base saturation percentage; SAR, sodium absorption ratio; Pav, available phosphorus; OC, Organic carbon; TN, total nitrogen; OC:TN, ratio of OC to TN.
TM, Teff (*Eragrostis tef* (Zucc) Trot)) mono-cropping; MM, Maize (*Zea mays L.*) mono-cropping; IC1; Cauliflower (*Brassica oleracea var. botrytis*) with maize intercropping; IC2, Red beet (*Beta Vulgaris*) with maize intercropping; R1, Cauliflower – teff - maize rotation; R2, Onion (*Allium cepa L.*) - maize - onion rotation; TG, Treated gully; UTG, Untreated gully; NF, Natural forest.

The highest soil EC was recorded from irrigated fields of IC2 (0.510 ds m^-1^) followed by R2 (0.390 ds m^-1^) whereas the lowest was found from rain-fed field of TM (0.057 ds m^-1^). However, there were non-significant differences in EC among many of the CLUS (e.g., MM, IC1, R1, R2, TG, UTG and NF) (Table 4). According to Landon [80], soil EC determined from the different CLUS is categorized as non-saline even though EC was found to be higher in the irrigation fields as compared to the other CLUS. The most likely reason for having low EC even in the irrigated fields could be attributed to the acceptable irrigation water quality, and irrigation method (furrow), and heavy rainfall during the summer season (June to early September) which may contribute for timely leaching of salts from the root zone. It is thus suggested to assess the salinity status of sub-surface soil layers of the irrigated fields in order to understand the extent of salt leached towards the lower soil horizons.

#### 3.2.2 Effects on cation exchange capacity and base forming cations

The highest CEC (50.3 cmol_c_ kg^-1^) and Ex K (1.43 cmol_c_ kg^-1^) were found from TG followed by NF (CEC = 49.8 cmol_c_ kg^-1^, Ex K =1.39 cmol_c_ kg^-1^) whereas the lowest CEC (11.4 cmol_c_ kg^-1^) and Ex K (0.204 cmol_c_ kg^-1^) were recorded from UTG. The significantly higher Ex Ca (25.5 cmol_c_ kg^-1^) and Ex Mg (15.1 cmol_c_ kg^-1^) were recorded from NF whereas the lowest Ex Ca (4.5 cmol_c_ kg^-1^) and Ex Mg (1.3 cmol_c_ kg^-1^) were observed from UTG (Table 4). The CEC, Ex Ca, Ex Mg, and Ex K recorded from MM were significantly higher than that of TM, IC1, IC2, R1, R2, and UTG. There were non-significant differences in CEC, exchangeable Ca, Mg, and K between the intercropping systems (IC1 and IC2), and also between crop rotation practices (R1 and R2) in the study sub-catchment. This indicates that crops used for intercropping and rotations have similar effect on CEC and soil exchangeable bases. However, CEC and those exchangeable bases recorded from irrigated fields with crop rotations were found to be higher than that of intercropping systems and rain-fed TM. Similarly, values of these soil properties were significantly higher in irrigated intercropping than rain-fed TM. Generally, CEC and exchangeable bases observed from NF and TG were categorized as very high; that of MM, IC1, IC2, R1 and R2 as high; from TM as medium; and UTG as a low rate as compared to the rates reported for African soils by Landon [80].

The highest Ex Na was found from IC2 (0.682 cmol_c_ kg^-1^) followed by IC2 (0.667 cmol_c_ kg^-1^). The lowest Ex Na was recorded from NF (0.030 cmol_c_ kg^-1^) and TM (0.040 cmol_c_ kg^-1^). The Ex Na recorded from TG was significantly higher than that of NF, TM, MM and UTG (Table 4), in which this could be attributed to the effects of long-term irrigation water drained to TG as it is located within the irrigation fields. According to Landon (1991), Ex Na observed from IC1, IC2, R1, R2 and TG were rated as medium; MM and UTG as low; and NF and TM as very low. However, the Ex Na from the irrigation fields was found to be near to the cut-off point for medium rate (0.7 cmol_c_ kg^-1^) which is regarded as potentially sodic, indicating that necessary soil and crop management practices should be taken to reduce or maintain Ex Na of the soil. In addition, the highest SBFC and BSP were found from NF and TG whereas the lowest was from UTG followed by TM (Table 4). According to the report by Landon [80] the BSP from NF and TG was rated as very high and that of UTG was rated as medium. According to the same author, the BSP of the remaining CLUS were categorized in the high rate. The highest ESP and SAR were recorded from IC1 followed by R2 and the lowest was from NF followed by TM (Table 4). However, all the CLUS showed ESP < 2, which is classified as low or non-sodic soils as reported in the rate for African soils by Landon (1991). Since the SAR is < 12 which is the cut-off point [58], the soil of the CLUS is categorized as non sodicity.

### 3.3 Effects on soil nutrients

The highest and statistically significant Pav was recorded from NF (23.9 mg kg^-1^) followed by TG (19.4 mg kg^-1^). However, the lowest Pav was found from UTG (1.4 mg kg^-1^) followed by TM (2.0 mg kg^-1^). The Pav contents among the intercropping and crop rotation practices under irrigation system were non-significantly differed. But Pav from MM was found to be significantly higher than the other cropping systems (Table 4). Soil Pav of NF, TG, and MM were rated as very high, high and medium, respectively, and the rest CLUS rated as very low in Pav as compared to the rates reported by Landon [80]). Soil management attention should thus be given to CLUS with very low soil Pav so that to improve Pav using appropriate practices and also maintain the Pav of potential land-use systems. Generally, this study result of Pav is consistent with previous reports elsewhere (e.g. Lemenin and Hanna [41]; Solomon et al. [83]; Nweke and Nnabude [84]; Flynn [85]) who stated that soil Pav variability has related to land-use type and soil management practices. For example, losses of Pav is higher in continuously cropped land as compared to forest land and other well managed land-use systems due to its fixation, removed with crop harvest, poor residual management and erosion processes ([84, 85].

The highest and statistically significant soil organic carbon (OC) was found from NF (4.98%) followed by TG (3.120%) while the lowest OC was from UTG (0.413%) followed by TM (0.643%). The optimal OC, i.e., between 3% < OC < 5 % as proposed by Craul [86], which indicates low risk of soil structural degradation is consistent with the values of NF and TG. The soil OC recorded from MM (1.45%) was significantly higher than that of intercropping (mean OC 1.31%) and crop rotation (mean OC 1.11%). The reason could be due to continuous application of manure or compost to MM than in the other arable fields and its residual effects on the sol system. The soil OC was higher in the intercropping fields than in the crop rotation which could be associated with the effect of legume crop (*Guizotia abyssinica* L.) intercropped with tef during the rain-fed crop season. Long-term studies on the benefit of manures, intercropping, and crop rotation have consistently reported for maintaining and increasing OC inputs into the soil and thereby impacts on soil properties [4, 14, 28]. However, even with intercropping, crop rotation and manure additions, continuous cropping and improperly intercropped and crop rotated fields resulted in a decline in OC. The rate and magnitude of the decline is affected by tillage, cropping system, management practices, and climate and soil conditions [14, 87].

According to Kay and Angers [88], irrespective of soil type and climatic condition if SOC contents are below 1% (e.g., TM, UTG), it may not be possible to obtain potential yields. Because SOC can impact on other physical, chemical and biological soil properties. In this study, SOC of TM and UTG showed very low; NF followed by TG showed very high and the SOC of the remaining CLUS (Table 4) are within the low rate as reported for African soils by Landon [80]. In agreement to the present finding other researchers elsewhere have reported that continuous cultivation depleted SOC and reduced soil quality compared to native vegetation (NF), regardless of the cropping system practiced (e.g., Reeves [87]; Bowman et al. [89]; Bremer et al. [90]).

The highest soil total nitrogen (TN) was found due to NF (0.541%) followed by TG (0.257%). The lowest TN was recorded from UTG (0.030%) followed by TM (0.067%). The soil TN from MM was significantly higher than that of IC1, IC2, R1 and R2 (Table 4), which could be attributed to the higher amount of manure applied to MM fields during each cropping season. TN from NF and TG rated as very high and high, respectively. The TN from the other CLUS rated as low except MM which is rated as medium and UTG as very low [80]. The present finding on OC and TN is consistent with other reports which have reported that SOC and TN content are not only affected by climate and terrain, but also by land-use and soil management practices. For example, agricultural intensification and repeated cultivation have resulted in a serious decrease in SOC and TN as compared to natural vegetation such as NF. The fact is that cultivation enhances decomposition of soil organic carbon, and physical removal of biomass as straw and grain harvest reduces its availability (e.g., Zhang et al. [61]; Wu et al. [63]; Nweke and Nnabude [84]; Johnson and Curtis [91]).

### 3.4 Effects on OC to TN ratio

The highest OC: TN was recorded from TG (13.7) followed by NF (12.9) and UTG (12.2) (Table 4), which could be associated with low oxidation (decomposition) rate of organic sources as compared to the inputs available in the study sub-catchment. Meaning, there were no soil and agronomic practices that enhanced decomposition of organic sources in these selected land-use systems. In addition, the soil in TG was water saturated almost for more than 8 months of the year, in which this could be slow down the decomposition of organic matter by limiting soil microbial activity [61, 63]. Similarly, long-term effects of irrigation practices can reduce microbial activity and thereby reduces organic matter decomposition which could be the reason for OC: TN to be slightly higher than 10 in the irrigation fields such as IC1, IC2, R1 and R2. The OC: TN of MM (11.4) was found to be higher than that of the fields under irrigation cropping system. The reason could be associated with the application of higher manure to MM field during the cropping season. It is also a fact that manure decomposes slowly during the irrigation cropping season. The lowest OC: TN was reported from TM (9.6), indicating that there is a higher organic matter mineralization. TM was the only CLUS which showed OC: TN below 10 (Table 4), indicating a balance in soil carbon and nitrogen nutrients [61, 92].

An optimum temperature and moisture conditions might be enhanced microbial activities to decompose organic sources in the fields such as TM [63]. However, the selected fields did receive little moisture from rainfall for about 8 months. The conventional tillage practice (cultivation) used in TM could also enhances organic sources to be decompose quickly [92, 93]. Tillage practice coupled with insufficient inputs of organic sources in the farming system of TM in a system that removes crop residue and absence of crop rotation resulted in a lower OC: TN [62, 94]. From agricultural production point of view literature showed that cropping and land-use systems with OC: TN < 10 is rated as good, 10.1-14 as medium and > 14 as poor soil systems [80]. However, such values are contrasted to the present pressures to reduce carbon emission to the atmosphere and sequestered carbon through maintaining higher soil OC: TN [62, 95, 96]. Hence, CLUS such as TG and NF showed better or conducive conditions for carbon sequestration as organic matter decomposition is slow down by limiting microbial activity as compared to cropping systems which accelerate the decomposition of organic sources. The OC: TN determined from TG and NF is lower than that of apple orchard (OC: TN of 15.4) reported for northern China [62, 96,]. However, the existing reports have not quantified the contribution of nitrogen deposition from the atmosphere to soil TN as this is the other factor that potentially affects the results of soil OC: TN of the different CLUS.

### 3.5 Effect on soil organic carbon (OC) and total nitrogen (TN) stocks

In the study sub-catchment, soil OC and TN stocks varied significantly among the majority of the cropping and land-use systems at 0-20 cm depth (Table 5). The highest stock of OC (113.6 Mg C ha^-1^) and TN (12.2 Mg C ha^-1^) were reported from NF. The soil OC stock from NF was slightly lower than that of reported for tropical forests (122 Mg C ha^_1^) by Prentice et al. (2001) in which such differences could be attributed to variability in N-fixing trees, soil factor and climatic conditions. The lowest stocks of soil OC (3.5 Mg C ha^-1^) and TN (0.25 Mg C ha^-1^) were found from UTG (Table 5). The soil OC and TN stocks of TG were significantly higher than the other CLUS, except that of NF. In line with the present results, Lal [25] has reported that soil organic matter (OM) can be greatly enhanced when degraded soils and ecosystems are restored, or converted to a restorative land-use or replanted to perennial vegetation (e.g., TG); and depleted OM in agricultural soils that use conventional tillage (e.g., TM). This could be the reason for OC to be used as an important indicator of both soil productivity and climate change mitigation interventions [25, 38].

**Table 5.** Mean SOC and TN stocks and reduction in stocks due to the cropping and land-use systems at 0- 20 cm soil depth in Dura sub-catchment, northern Ethiopia.

| Cropping and land-use system | SOC stock (Mg C ha<sup>-1</sup>) | SOC stock reduction (%) | TN stock (Mg N ha<sup>-1</sup>) | TN stock reduction (%) |
| --- | --- | --- | --- | --- |
| TM | 10.6e | 90.7 | 1.09e | 91.0 |
| MM | 28.5c | 74.9 | 2.67c | 78.0 |
| IC1 | 23.4d | 79.4 | 2.05d | 83.1 |
| IC2 | 19.7d | 82.7 | 1.68de | 86.2 |
| R1 | 16.7de | 85.3 | 1.60de | 86.8 |
| R2 | 17.1de | 84.9 | 1.28de | 89.5 |
| TG | 49.4b | 56.5 | 3.65b | 70.0 |
| UTG | 3.50f | 96.9 | 0.25f | 97.9 |
| NF | 113.6a | 0.00 | 12.2a | 0.00 |
| Mean | 31.0 | 72.4 | 2.92 | 75.8 |

TM, Teff (*Eragrostis tef* (Zucc) Trot)) mono-cropping; MM, Maize (*Zea mays L.*) mono-cropping; IC1, Cauliflower (Brassica oleracea var. botrytis) with maize intercropping; IC2, Red beet (*Beta Vulgaris*) with maize intercropping; R1, Cauliflower – teff - maize rotation; R2, Onion (*Allium cepa L.*)- maize - onion rotation; TG, Treated gully; UTG, Untreated gully; NF, Natural forest land system.

The soil OC and TN stocks estimated from MM were showed significantly higher than all the other CLUS, except that of NF and TG. This could be associated with the seasonal application of organic inputs on MM fields. The soil OC and TN stocked in the intercropped fields were higher than that of rotation and TM which might be associated with the relatively effectiveness of the intercropping system for improving soil OC sequestration. Overall, such OC and TN stock variability is attributed to decrease in soil bulk density values significantly across the CLUS. The present results of OC and TN stocks variability across the CLUS is agreed to the previous reports of Bird et al. [97] and Saiz et al. [98] who have reported variability in soil stocks attributed to different factors across land-use and land management practices. For example, practices in NF and TG (Table 1) that reduce soil OM mineralization and erosion, and increase OM inputs to the soil could improve soil OC and TN stocks.

As compared to the reference land-use system (NF), soil OC stock due to UTG and TM was reduced by 97 and 91%, respectively. Similarly, TN stock was reduced due to UTG and TM by 98 and 91%, respectively. The mean reduction of OC stocks by crop rotation, intercropping, MM, and TG as compared to the NF were found to be 85, 81, 75, and 57%, respectively. The mean TN stock reduced due to crop rotation, intercropping, MM, and TG as compared to NF was calculated as 88, 85, 78, and 70%, respectively. This study results thus indicated that the SOC and TN stocks are drastically reduced in most of the CLUS, with the highest reduction from UTG followed by TM. Losses of OC and TN from the soil system could increase the amount of carbon and nitrogen gasses in the atmosphere at global-scale [38, 99]. The TN stock reduction is slightly higher than the SOC stock across all the CLUS, indicating that understanding the reason for more soil nitrogen depletion should merit further investigation. Generally, the soil OC stock reduced in this study is reported higher than the previous reports from Africa (e.g., Amudson [99]) who reported that cultivation reduces the original OC content by up-to 30%. In the study catchment, such variability could be attributed to the duration and management practices of the CLUS as arable lands have been cultivated for more than 100 years. In addition, increasing soil OC stock is a major challenge in dry areas where vegetation growth is hampered by climatically harsh conditions such as low availability of water and nutrients, removal for fodder and fuelwood [38].

### 3.6 Determinants of soil prosperities variability using PCA

The bivariate correlation analysis among many soil properties determined from the CLUS correlated (r) at r > 0.70 which qualitatively described as moderate to extremely strong correlation (data not shown). Such values of r indicate that there are multicollinearity effects among the soil properties. The effect of multicollinearity was handled using principal component analysis (PCA) that grouped soil properties into five principal components (PCs) (Table 6). The eigenvalue of PC1, PC2, PC3, PC4 and PC5 are 8.50, 6.48, 2.24, 1.61 and 1.13, respectively. The five PCs that received eigenvalues >1 explain largely the variability of the soil properties among the CLUS [65]. The variances explained by PC1, PC2, PC3, PC4 and PC5 are 30.65, 24.32, 17.29, 7.90 and 6.63, respectively. Such variance values are in line with the report of Brejda et al. [65] who sated that PCs that receive at least 5% variance explains the best variability of a factor component. The soil properties included in the first five PCs explain for 86.8% of the soil quality variability among the CLUS. The communalities of the extracted five PCs explained by each soil property ranges from 74-96% (Table 6). A high communality variable shows that a high portion of variance was explained the variable and therefore, it gets a higher preference to a low communality [100].

**Table 6.**
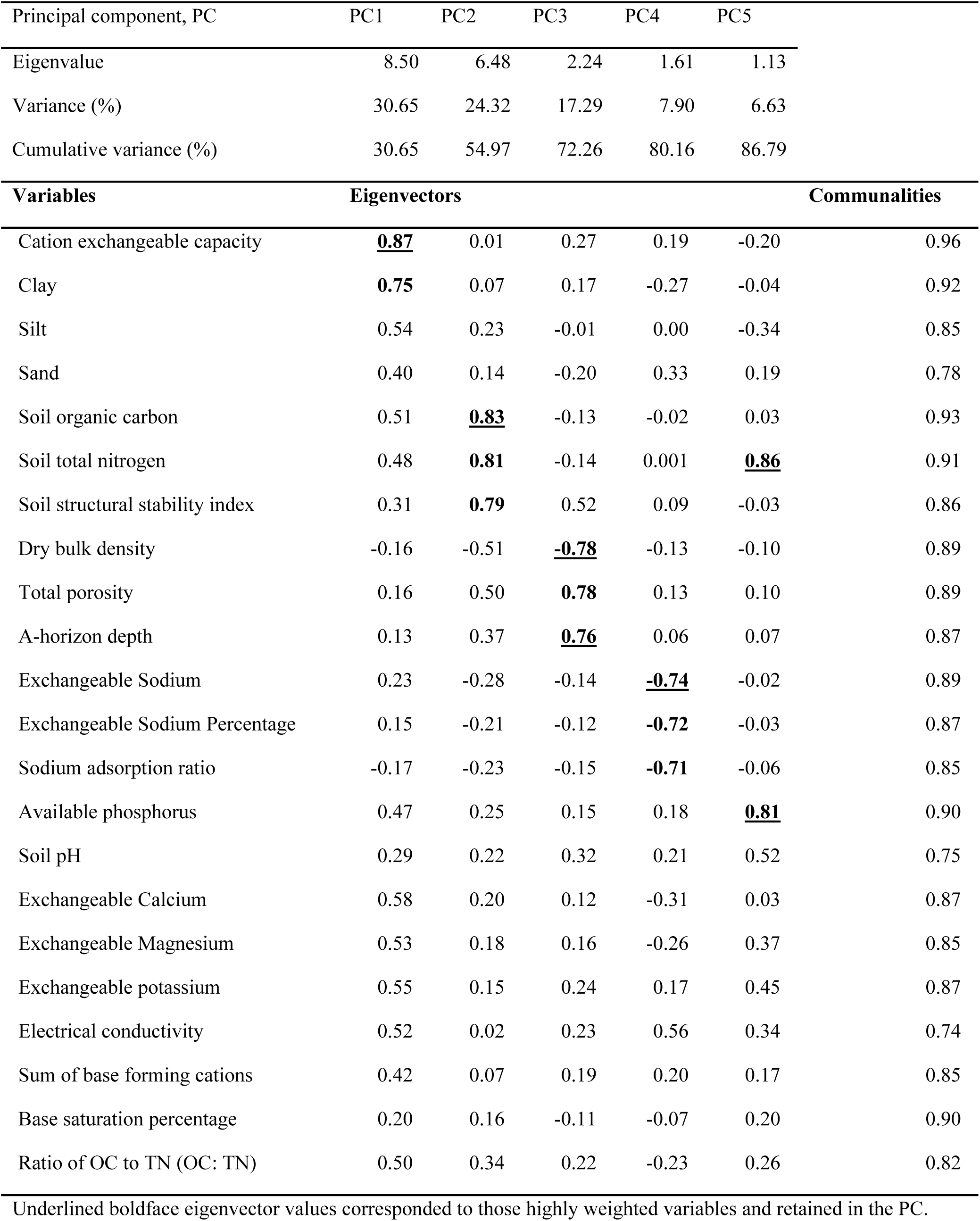
Results of principal component analysis of soil properties responses to the cropping and land-use systems in the Dura sub-catchment, northern Ethiopia.

| Principal component, PC | PC1 | PC2 | PC3 | PC4 | PC5 |  |
| --- | --- | --- | --- | --- | --- | --- |
| Eigenvalue | 8.50 | 6.48 | 2.24 | 1.61 | 1.13 |  |
| Variance (%) | 30.65 | 24.32 | 17.29 | 7.90 | 6.63 |  |
| Cumulative variance (%) | 30.65 | 54.97 | 72.26 | 80.16 | 86.79 |  |
| Variables | Eigenvectors |  |  |  |  | Communalities |
| Cation exchangeable capacity | **<u>0.87</u>** | 0.01 | 0.27 | 0.19 | -0.20 | 0.96 |
| Clay | **0.75** | 0.07 | 0.17 | -0.27 | -0.04 | 0.92 |
| Silt | 0.54 | 0.23 | -0.01 | 0.00 | -0.34 | 0.85 |
| Sand | 0.40 | 0.14 | -0.20 | 0.33 | 0.19 | 0.78 |
| Soil organic carbon | 0.51 | **<u>0.83</u>** | -0.13 | -0.02 | 0.03 | 0.93 |
| Soil total nitrogen | 0.48 | **0.81** | -0.14 | 0.001 | **<u>0.86</u>** | 0.91 |
| Soil structural stability index | 0.31 | **0.79** | 0.52 | 0.09 | -0.03 | 0.86 |
| Dry bulk density | -0.16 | -0.51 | **<u>-0.78</u>** | -0.13 | -0.10 | 0.89 |
| Total porosity | 0.16 | 0.50 | **0.78** | 0.13 | 0.10 | 0.89 |
| A-horizon depth | 0.13 | 0.37 | **<u>0.76</u>** | 0.06 | 0.07 | 0.87 |
| Exchangeable Sodium | 0.23 | -0.28 | -0.14 | **<u>-0.74</u>** | -0.02 | 0.89 |
| Exchangeable Sodium Percentage | 0.15 | -0.21 | -0.12 | **-0.72** | -0.03 | 0.87 |
| Sodium adsorption ratio | -0.17 | -0.23 | -0.15 | **-0.71** | -0.06 | 0.85 |
| Available phosphorus | 0.47 | 0.25 | 0.15 | 0.18 | **<u>0.81</u>** | 0.90 |
| Soil pH | 0.29 | 0.22 | 0.32 | 0.21 | 0.52 | 0.75 |
| Exchangeable Calcium | 0.58 | 0.20 | 0.12 | -0.31 | 0.03 | 0.87 |
| Exchangeable Magnesium | 0.53 | 0.18 | 0.16 | -0.26 | 0.37 | 0.85 |
| Exchangeable potassium | 0.55 | 0.15 | 0.24 | 0.17 | 0.45 | 0.87 |
| Electrical conductivity | 0.52 | 0.02 | 0.23 | 0.56 | 0.34 | 0.74 |
| Sum of base forming cations | 0.42 | 0.07 | 0.19 | 0.20 | 0.17 | 0.85 |
| Base saturation percentage | 0.20 | 0.16 | -0.11 | -0.07 | 0.20 | 0.90 |
| Ratio of OC to TN (OC: TN) | 0.50 | 0.34 | 0.22 | -0.23 | 0.26 | 0.82 |

Underlined boldface eigenvector values corresponded to those highly weighted variables and retained in the PC.

The highly loading variables in PC1 were CEC and clay content (Table 6). Since the correlation coefficient between CEC and clay was 0.86 which is higher than the cut-off point (0.60,), communality was considered to select the parameter to be retained in PC1. As a result, CEC was retained in PC1 because the loading (0.87) and communality (0.95) of CEC were higher than that of clay. The first PC is thus termed as ‘*cation exchange capacity, CEC factor’*. Similarly, the highly loading variables under PC2 were SOC, TN and soil structural stability index (SSSI). However, the partial correlation analysis results indicated strong correlations (r > 0.80) among these variables. Considering the higher partial correlation coefficient, loading value and communality, SOC was retained in PC2. The content of SOC influences directly the value of the other highly loading variables (TN and SSSI) [82, 101, 102]. The implication these reports is that the contribution of TN and SSSI to PC2 is explained using SOC. As a result, PC2 is termed as the ‘*organic matter factor’*. The highly loaded variables in PC3 included dry soil bulk density (DBD), porosity and A-horizon depth (A_h). The partial correlation analysis between DBD and porosity showed at r = 0.85. Since the loading value and communality of DBD was slightly higher than that of porosity, DBD was retained in PC3. A-horizon depth was also retained in PC3 as the correlation coefficient with the other high loading variables showed less than the cut-off point (< 0.60). Thus, PC3 is termed as *‘soil physical property* factor’. The highly loaded variable in PC4 included Ex Na, ESP and SAR. The partial correlation values among these variables showed strong correlation (r > 0.88). Considering the higher correlation coefficient, loading weight and communality values, Ex Na was retained in PC4 and the rest variables were excluded from PC4. As a result, PC4 was termed as the ‘*sodicity factor’*. Likewise, the highly loaded variables in PC5 are TN and Pav (Table 6). Since the correlation between TN and Pav is below 0.6, both variables were retained in PC5. The loading coefficient of TN in PC5 (0.86) is higher than in PC2 (0.81) which is another reason to retain TN in PC5. Nitrogen and phosphorus are commonly reported as the most crop limiting soil nutrients in developing countries. So, considering these parameters in PC5 is important while assessing soil degradation using soil properties as an indicator among the CLUS. Thus, PC5 is termed as the ‘*soil macro-nutrient factor’*.

The seven PCA chosen final soil properties that better explain (determinant) of soil quality variability among the CLUS were CEC, SOC, DBD, A_h, Ex Na, TN, and Pav. Future assessment and monitoring of the effects of CLUS on soil quality is suggested to depend on these seven soil properties which are sensitive to disturbances related to land-use and management practices. This can reduce wastage of resources (e.g., cost, time, labor) while analyzing the entire datasets and also gives rapid response for effective assessment and decision-making. Generally, among the seven PCA chosen soil properties of this study, five (CEC, SOC, TN, phosphorus, DBD) parameters were similarly selected by Tesfahunegn et al. [82] who has reported from exclusively rain-fed land-use and soil management practices in northern Ethiopia. The choice of SOC in the PC factor as determinant soil property for soil quality variability across the CLUS is consistent with other reports (e.g., Larson Pierce [103]; Shukla et al. [104]) who have reported that soil organic matter (SOM) is among the most powerful soil properties to influence many soil conditions. For example, Larson and Pierce [103] have reported that SOM improves the soil to accept, hold, and release nutrients, water and other soil chemical ingredients to plants, recharge surface water to groundwater; support root growth through soil structure stability, maintain soil biotic habitat; and resist soil degradation. The selection of DBD using the PCA could also confirm the basic principle of the soil to restrict water flow and plant root growth when DBD increases [105, 106].

## Conclusions

This study revealed that soil properties and nutrient stocks affected significantly by the cropping and land-use systems (CLUS) in the study sub-catchment. The natural forest (NF) followed by the treated gully (TG) were attempted to maintain the sustainability of the soil properties as compared to the other CLUS. All the soil chemical properties except exchangeable sodium (Ex Na) showed the highest values in NF and TG. Similarly, NF followed by TG showed a lower dry bulk density than the other CLUS. Conversely, the soil properties and nutrient stocks determined from the untreated gully (UTG) followed by tef mono-cropping (TM) showed a seriously degraded soil quality indicators. The soil properties status determined from maize mono-cropping (MM) is by far better than that of irrigated intercropping and crop rotation fields. The highest Ex Na was reported from the irrigation fields, particularly from the intercropping systems in which this demands serious attention to decrease and maintain soil sodium status. However, most of the other soil properties and nutrient stocks (SOC and TN) showed a better improvement in intercropping than crop rotation fields. In this study, the soil properties values generally showed an early warning about the severity of soil physical and nutrient degradation in some of the CLUS such as UTG and TM followed by the irrigated fields. In the study area, the CLUS having a better soil prosperities and nutrient stocks were reported in descending order as NF, TG and MM. It is thus suggested that implementation of appropriate intercropping and crop rotation systems (e.g., use of legume crops and trees) and best soil management practices to improve and maintain the soil quality of the study sub-catchment conditions. For further monitoring of the effects of the CLUS on soil quality in the study sub-catchment conditions, attention should be given to the PCA selected variables (CEC, SOC, DBD, A_h, Ex Na, TN, and Pav). These seven variables are explained for 87% of the soil quality variability due to the CLUS. Such selections of few relevant soil parameters are helpful as a means to assess soil properties that leads to quick monitoring (rapid and inexpensive) and effective decision-making against the effects of each CLUS on soil properties and nutrient stocks in the study sub-catchment conditions.

## Abbreviations

OC: organic carbon;
TN: total nitrogen;
EC: electrical conductivity;
CEC: cation exchangeable capacity;
Ex Na, Ex Ca, and Ex Mg: exchangeable sodium, calcium, magnesium, respectively;
Pav: available phosphorus;
A_h: depth of A-horizon;
PCA: principal component analysis.

## Data availability

The data used to support the findings of this study are available from the corresponding author upon request.

## Acknowlegements

This research was financial supported by University of Aksum (Ethiopia) under the terms of grant no. AKU/IG/RCSD/1092/07. The author gratefully acknowledged the financial support provided by Aksum University to conduct this study. The author also highly appreciated the farmers and development agents who involved in the identification and characterization of the different cropping and land-use systems. The support provided during the soil sample collection by Mr. Kahsu Kidane (Development Agent) is highly appreciated. The assistance offered by the village administration and development agents during all the discussions and data collection processes are also highly acknowledged.

